# A Single Dose of Inactivated Influenza Virus Vaccine Expressing COBRA Hemagglutinin Elicits Broadly-Reactive and Long-Lasting Protection

**DOI:** 10.1101/2024.05.13.593905

**Authors:** Hua Shi, Xiaojian Zhang, Ted M. Ross

**Affiliations:** Center for Vaccines and Immunology, University of Georgia, Athens, GA, USA; Florida Research and Innovation Center, Cleveland Clinic, Port Saint Lucie, FL, USA; Department of Infectious Diseases, University of Georgia, Athens, GA, USA; Department of Infection Biology, Lehner Research Institute, Cleveland Clinic, Cleveland, OH, USA

**Keywords:** inactivated influenza vaccine, pre-immunity, COBRA, transmission, adjuvants

## Abstract

Influenza virus infections present a pervasive global health concern resulting in millions of hospitalizations and thousands of fatalities annually. To address the influenza antigenic variation, the computationally optimized broadly reactive antigen (COBRA) methodology was used to design influenza hemagglutinin (HA) or neuraminidase (NA) for universal influenza vaccine candidates. In this study, whole inactivated virus (WIV) or split inactivated virus (SIV) vaccine formulations expressing either the H1 COBRA HA or H3 COBRA HA were formulated with or without an adjuvant and tested in ferrets with pre-existing anti-influenza immunity. A single dose of the COBRA-WIV vaccine elicited a robust and broadly reactive antibody response against H1N1 and H3N2 influenza viruses. In contrast, the COBRA-SIV elicited antibodies that recognized fewer viruses, but with R-DOATP, its specificity was expanded. Vaccinated ferrets were protected against morbidity and mortality following challenge with A/California/07/2009 at 14 weeks post-vaccination with reduced viral shedding post-infection compared to the naïve mock-vaccinated ferrets. However, the COBRA-IIVs did not block the viral transmission to naïve ferrets. The contact infection induced less severe disease and delayed viral shedding than direct infection. Overall, the COBRA HA WIV or the COBRA HA SIV plus R-DOTAP elicited broadly reactive antibodies with long-term protection against viral challenge and reduced viral transmission following a single dose of vaccine in ferrets pre-immune to historical H1N1 and H3N2 influenza viruses.

**IMPORTANCE:** A next-generation influenza virus vaccine aims to provide broadly reactive protection against various drifted influenza strains. COBRA HA-based vaccines elicit broadly-reactive antibodies following two or three vaccinations. However, people are administered a single vaccination of current commercial influenza virus vaccines. In this study, ferrets with pre-existing anti-influenza immunity were administered a single shot COBRA-WIV that elicited long-lasting, broadly-reactive antibodies that protected ferrets against pdmH1N1 viral challenge. This is the first study describing the immune responses elicited by COBRA-IIV vaccines in ferrets that protected ferrets 14 weeks post-vaccination.

## Introduction

Influenza viruses are a worldwide public health threat that causes respiratory tract infections and, in more severe cases, pneumonia and death (1). Each year, influenza virus infections are responsible for over 5 million hospitalizations for adults and 10.1 million influenza virus-associated acute lower respiratory infections with 34,800 reported deaths for children under 5 years old globally (2, 3). The Centers for Disease Control and Prevention (CDC) recommends prophylactic vaccination as the most effective method for preventing influenza virus breakouts (https://www.cdc.gov/flu/season/faq-flu-season-2023-2024.htm). In recent flu seasons, the most commonly used seasonal influenza vaccine is the split inactivated vaccine (SIV) consisting of one H1N1 and one H3N2 influenza A virus strain, and two influenza B virus strains selected before the flu season (4). However, the vaccine effectiveness (VE) was 40%-50% for the recent vaccines (5, 6), and could be much lower (only 19% for the 2014-2015 vaccine) when a mismatch between the vaccine strains and circulating strains occurs (7). Developing a universal influenza vaccine (UIV) that can provide broad protection against multiple strains for all populations is an unmet goal for the next-generation influenza vaccines.

Hemagglutinin (HA) binds to the sialic acid and facilitates the entry of virions (8), serving as the primary target for the vaccine design (9–11). To design the HA structure that can elicit broadly reactive antibodies against various influenza strains, the computationally optimized broadly reactive antigen (COBRA) technology was developed by using multiple layered consensus building (12). COBRA H1, H2, and H5 have been designed in recombinant HA (rHA) or virus-like-particle (VPL) formats, and their effectiveness in eliciting antibodies with a broad spectrum has been confirmed in mouse and ferret models (13–16).

In this study, the inactivated influenza vaccine (IIV) expressing COBRA HA was assessed as a UIV candidate. The whole inactivated influenza vaccine (WIV) was first used in humans to prevent influenza viral infection in 1940 (17), but later the SIV replaced the WIV in the U.S. market for its improved safety (18, 19). In a previous study, the bivalent COBRA-SIV carrying both COBRA H1 and H3 was designed and tested in naïve ferrets. The results showed that the COBRA-SIV elicited broader or similar antibody reactivities compared to the wild-type (WT) SIV (20). However, the COBRA-SIV failed to elicit protective antibody levels against all strains in the panels, which could be because of the reduced antigenicity of SIV. Later, the WIV or SIV expressing an H1 (Y2) or H3 (J4) COBRA HA antigen was designed and tested in mice models for a more comprehensive evaluation of the COBRA HA-based IIVs (H. Shi, X. Zhang, P. Ge, V. Meliopoulos, P. Freiden, B. Livingston, S. Schultz-Cherry, T.M. Ross, submitted for publication). The results showed that the addition of an adjuvant enhanced the long-term protection elicited by both COBRA-WIV and COBRA-SIV and the pre-immunity further expanded the antibody activity spectrum (H. Shi, X. Zhang, P. Ge, V. Meliopoulos, P. Freiden, B. Livingston, S. Schultz-Cherry, T.M. Ross, submitted for publication).

To further evaluate the effectiveness of COBRA-IIVs in eliciting antibodies with broad spectrum under different formulations and restricting viral transmission, herein, the ferret transmission study is designed. The adjuvants are commonly used in vaccines to improve the VE, for example, the adjuvanted influenza vaccine was approved for the elderly in the U.S. (21). Therefore, in this study, two adjuvants, AddaVax and R-DOTAP, served as active competitors for optimizing the design of COBRA-IIVs. AddaVax, as an equivalent of the commercial influenza vaccine adjuvant MF59, can stimulate a balanced Th1/Th2 activation (22). The novel cationic nanoparticle R-DOTAP can stimulate cellular immunity, especially CD8T cells, to eliminate intracellular pathogens (23, 24). In the previous study, the primary vaccination of COBRA-IIVs successfully recalled memory B cells in the pre-immune mice (H. Shi, X. Zhang, P. Ge, V. Meliopoulos, P. Freiden, B. Livingston, S. Schultz-Cherry, T.M. Ross, submitted for publication), which indicates that a single dose of COBRA-IIV is sufficient to elicit immune memory. To better evaluate the effectiveness of a single dose of a COBRA-IIV vaccine, pre-immune ferrets were vaccinated with either COBRA-WIV or COBRA-SIV vaccine followed by an H1N1 influenza virus challenge 14 weeks later. Ferrets were assessed for high titer antibodies with HAI activity, protection against direct infection, and prevention of viral shedding and transmission.

## Results

### COBRA-IIVs elicited broadly reactive antibodies against multiple H1N1 and H3N2 strains

This study assessed the effectiveness of the COBRA-IIV vaccines to elicit antibodies with broadly reactive HAI activity following a single vaccination in ferrets with pre-existing immunity to historical H1N1 and H3N2 influenza strains (Figure 1). Eight weeks following infection with historical influenza strains, ferrets were vaccinated with either WIV or SIV vaccine expressing COBRA HA antigens formulated with R-DOTAP or AddaVax adjuvant or nonadjuvanted (Table 1). Two weeks following vaccination, ferrets had antibodies with HAI activity (Figure 2). The mock vaccinated ferrets had no detectable HAI activity, therefore, only HAI activity is shown for pre-immune, vaccinated ferrets. Ferrets with pre-existing anti-influenza virus immunity had antibodies with HAI activity (average titer ≥1:40) at 2 weeks following COBRA-WIV vaccination against 50% of the H1N1 influenza virus strains and 100% of the H3N2 influenza virus strains (Figure 2A). In contrast, ferrets vaccinated with these same vaccines plus R-DOTAP had similar results (Figure 2B), whereas ferrets vaccinated with these same vaccines plus AddaVax adjuvant had antibodies with HAI activity against 75% of the H1N1 influenza virus strains, but only 43% of the H3N2 influenza virus strains (Figure 2C). Ferrets vaccinated with COBRA-SIV alone or with adjuvant had antibodies with HAI activity that recognized fewer H1N1 or H3N2 influenza viruses in the panel compared to their COBRA-WIV counterparts (Figure 2D-F). HAI antibody titers against CA/09, the influenza challenge strain, were assessed for 14 weeks post-vaccination (Figure S1). Regardless of the vaccine administered, ferrets had a four-fold decline in HAI activity against CA/09-specific at week 14 compared to week 2 post-vaccination.

**Figure 1.**
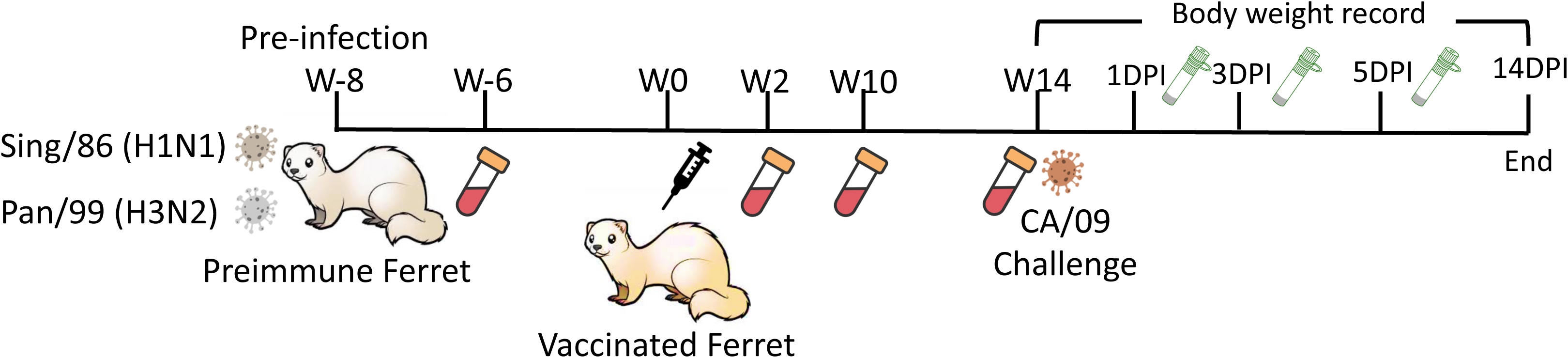
Experimental timeline for vaccination and direct challenge. 36 naïve ferrets were pre-infected with Sing/86 and Pan99 (5*10^5^PFU/virus/ferret) 8 weeks prior to the vaccination. Sera samples were collected 2 weeks post-pre-infection. At week 0, ferrets were evenly divided into 6 groups receiving vaccines named: COBRA-WIV, COBRA-WIV plus AddaVax, COBRA-WIV plus R-DOTAP, COBRA-SIV, COBRA-SIV plus AddaVax, and COBRA-SIV plus R-DOTAP. The detailed information of each vaccine’s formulation is shown in Table 1. Sera samples were collected on week 2, 10, and 14 to measure the antibody levels. Ferrets were challenged with CA/09 (1*10^6^PFU/ferret) at week 14, with 6 naïve mock-vaccinated ferrets serving as the mock control. Nasal washes were collected at 1DPI, 3DPI, and 5DPI to monitor the viral shedding. Body weight and clinical signs were closely recorded daily until 14DPI. DPI: day post-infection.

**Figure 2.**
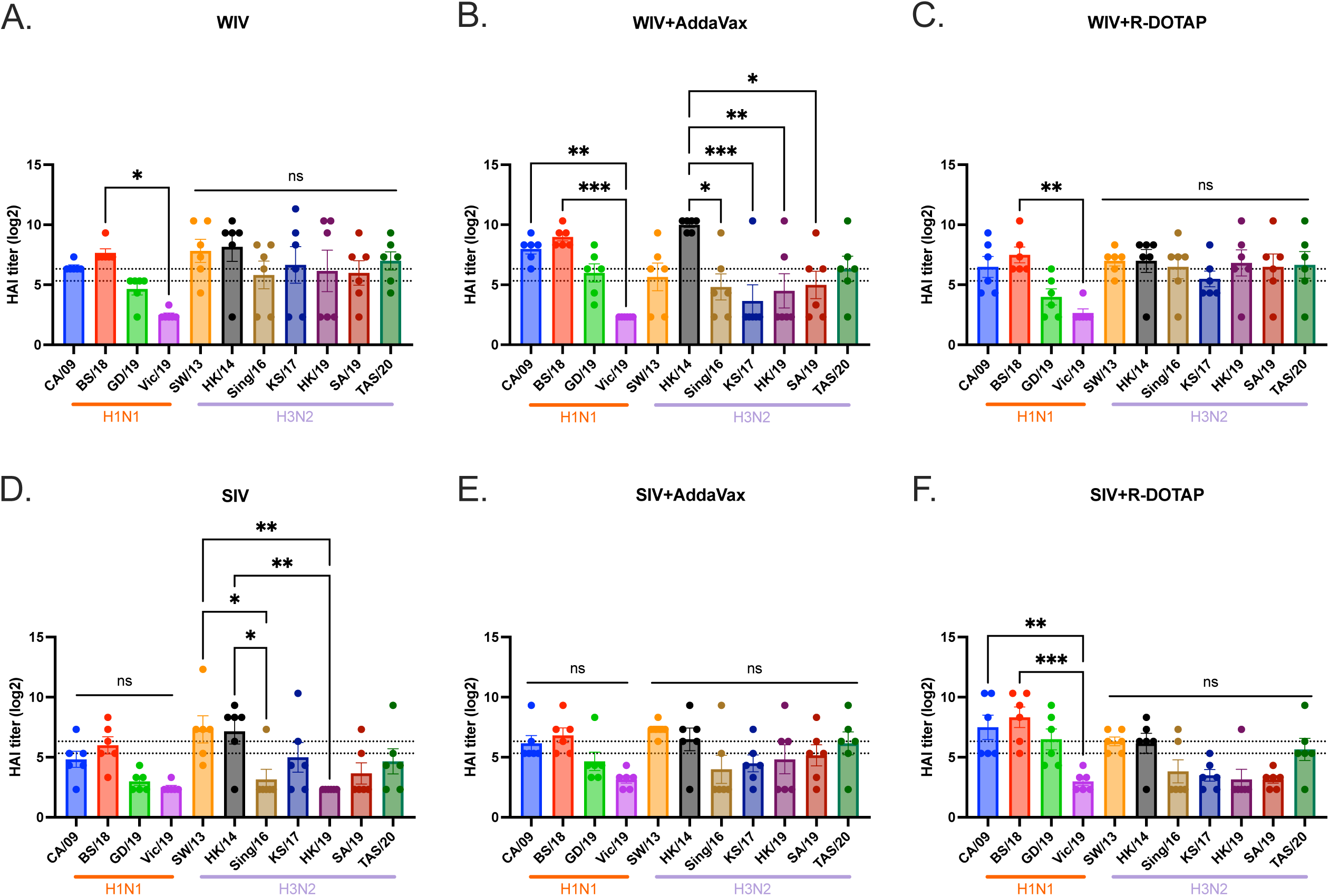
HAI titers against H1N1 and H3N2 panels for sera collected 2 weeks post prime vaccination. The HAI titers elicited by COBRA-WIV (A), COBRA-WIV plus AddaVax (B), COBRA-WIV plus R-DOTAP (C), COBRA-SIV (D), COBRA-SIV plus AddaVax (E), and COBRA-SIV plus R-DOTAP (F) were measured. The X-axis represents different virus strains. The red bar indicates the H1N1 strains and the purple bar indicates the H3N2 strains. The Y-axis represents the Log2 HAI titers with absolute mean values ± SEM. The lower dotted line indicates the HAI titer of 1:40, and the upper dotted line indicates 1:80. A P value of less than 0.05 was defined as statistically significant (*, P < 0.05; **, P < 0.01; ***, P < 0.001; ****, P < 0.0001).

**Table 1.** Formulation information for the COBRA-IIVs.

| **Group** | **Vaccine** | **Dose** | **Adjuvant** |
| --- | --- | --- | --- |
| 1 | COBRA Y2+J4 WIV | 7.5ug/HA/ferret | N/A |
| 2 | COBRA Y2+J4 WIV | 7.5ug/HA/ferret | AddaVax |
| 3 | COBRA Y2+J4 WIV | 7.5ug/HA/ferret | R-DOTAP |
| 4 | COBRA Y2+J4 SIV | 7.5ug/HA/ferret | N/A |
| 5 | COBRA Y2+J4 SIV | 7.5ug/HA/ferret | AddaVax |
| 6 | COBRA Y2+J4 SIV | 7.5ug/HA/ferret | R-DOTAP |

### COBRA-IIV vaccines elicit broad neutralizing antibodies against H1N1 and H3N2 panels

To assess the ability of the vaccine-elicited sera to neutralize virus infection, a microneutralization assay was performed (Figure 3).

**Figure 3.** The breadth of neutralizing antibodies elicited by COBRA-IIVs 2 weeks post prime vaccination. The MNA titers elicited by CORBA-IIVs were measured against H1N1 strains CA/09 (A) and Vic/19 (B), and H3N2 strains Sing/16 (C), KS/17 (D), HK/19 (E), SA/19 (F), and TAS/20 (G). The X-axis represents different vaccines. The Y-axis represents the Log2 MNA endpoint with absolute mean values ± SEM. MNA: microneutralization assay. MNA experiments ongoing

and the results are…

Experiment ongoing

### CORBA-IIV vaccines stimulate the IgG isotype switch and HA stem-binding antibodies

Anti-HA IgG antibody titers were measured 2 weeks post-vaccination (Figure 4). On average, all ferrets vaccinated with the COBRA-WIV vaccine with or without adjuvant had statistically similar anti-H1 HA antibody titers (Figure 4A). Ferrets vaccinated with COBRA-SIV vaccines plus R-DOTAP doubled anti-HA antibody titers compared to ferrets vaccinated with COBRA-SIV vaccine only. All ferrets had statistically similar anti-H3 HA antibodies following vaccination, regardless of the vaccine or adjuvant administered (Figure 4B). Ferrets vaccinated with the COBRA-SIV vaccine plus R-DOTAP had more antibodies directed to the group 1 HA stem region compared to ferrets vaccinated with the COBRA-SIV vaccine only or with the COBRA-SIV vaccine plus AddaVax (Figure 4C). In contrast, ferrets vaccinated with the COBRA-WIV vaccine only had significantly higher to the group 1 stem HA compared to ferrets vaccinated with these same vaccines with either adjuvant. Additionally, the COBRA-WIV vaccine only stimulated significantly more group 1 stem-binding antibodies than the COBRA-SIV vaccine only. Overall, all ferrets had similar group 2 anti-HA stem antibodies following vaccination with either vaccine with or without adjuvants (Fig 4D). Taken together, one vaccination of either COBRA-IIV vaccine elicits anti-HA head and stem antibodies following vaccination.

**Figure 4.**
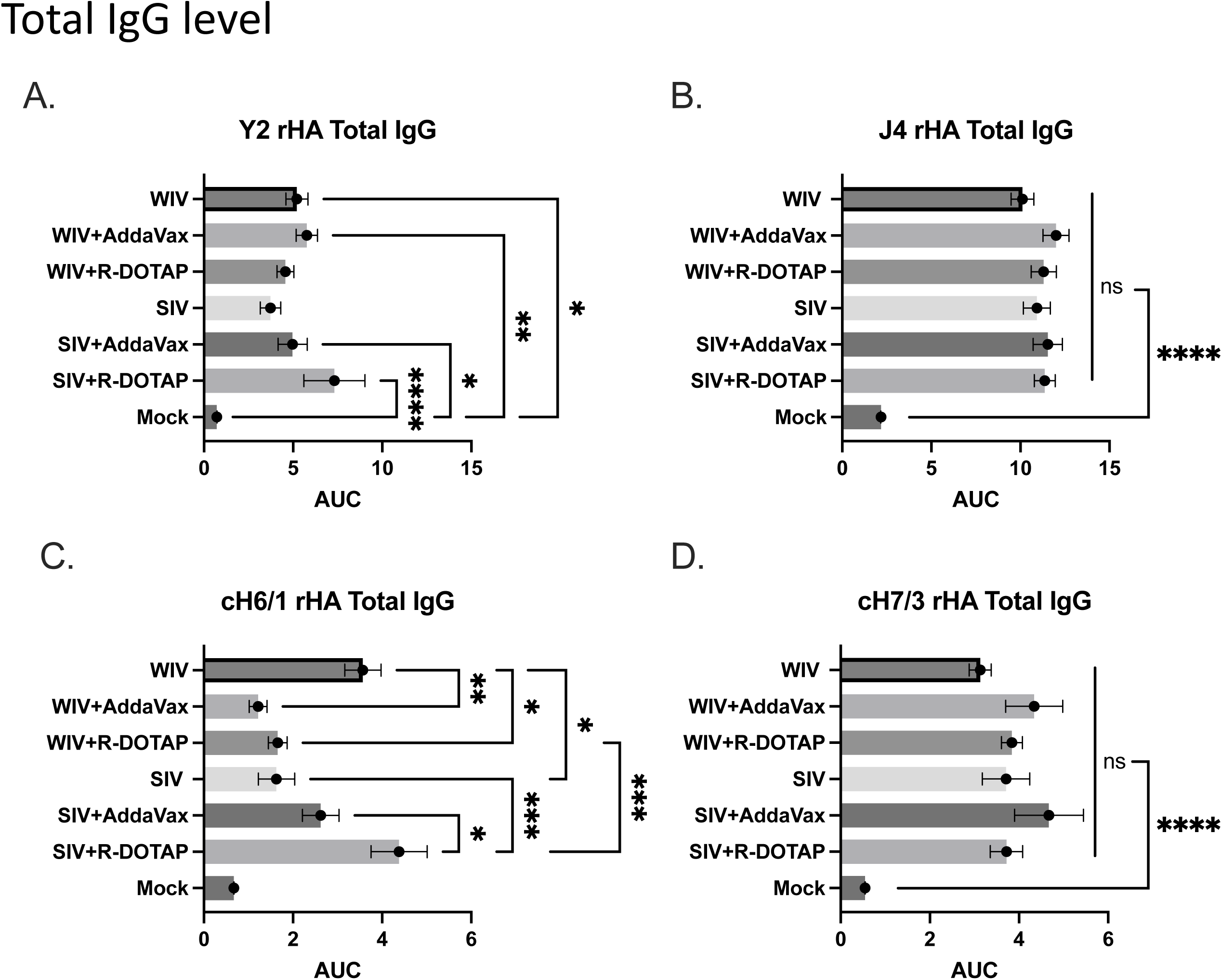
IgG levels stimulated by COBRA-IIVs 2 weeks post prime vaccination. A. Y2-specific total IgG level. B. J4-specific total IgG level. C. H1 HA stem-binding IgG level. D. H3 HA stem-binding IgG level. The X-axis represents different vaccines. The Y-axis represents the Log2 MNA endpoint with absolute mean values ± SEM. The X-axis represents the area under curve (AUC) obtained OD141 values from 3-fold serially diluted sera plus SEM. The Y-axis represents different vaccines. cH6/1: chimeric rHA with H6 head from H6 A/Mallard/Sweden/81/2002 and stalk from A/California/07/2009. cH7/3: chimeric rHA with H7 head from A/Anhui/1/2013 and H3 stalk from A/Texas/50/2012. A P value of less than 0.05 was defined as statistically significant (*, P < 0.05; **, P < 0.01; ***, P < 0.001; ****, P < 0.0001).

### COBRA-IIVs mitigated clinical signs and viral shedding in pre-immune ferrets after CA/09 exposure

At 14 weeks post-vaccination, vaccinated ferrets were challenged via intranasal inoculation with the H1N1 influenza virus, CA/09 (1*10^6^PFU/ferret). Unvaccinated ferrets rapidly lost weight following the challenge losing ∼15% of their original body weight by day 7 post-infection (Figure 5A) with a third of the ferrets succumbing to infection (Figure 5B). All ferrets vaccinated with COBRA-WIV vaccines, with or without adjuvant, lost between 5-10% body weight in the first 3-4 days post-infection and then, began returning to their original weight over the 14 days of observation (Figure 5A). All COBRA-WIV vaccinated ferrets survived challenge (Figure 5B). Similar results were observed in ferrets vaccinated with COBRA-SIV vaccine with or without adjuvant, except a third of the ferrets vaccinated with COBRA-SIV only vaccines died from challenge (Figure 5B). During the 14-day period, mild clinical symptoms were observed in all ferrets, such as sneezing and nasal discharge and several ferrets in the mock control group were lethargic (Figure S2). Mock vaccinated ferrets had high viral nasal wash titers (1*10^6^ to 1*10^7^ PFU/ml) at 1-day post-infection (Figure 6). Ferrets vaccinated with any vaccine or adjuvant had similar viral nasal wash titers at 1-day post-infection. Influenza virus was detected from the nasal washes of ferrets following the CA/09 challenge (Figure 6). Viral nasal wash titers dropped a log by day 3 post-infection in mock vaccinated mice (Figure 6) and dropped an additional log to 1*10^4^ PFU/ml at day 5 post-infection. In contrast, vaccinated ferrets had viral nasal wash titers that ranged on average from 1*10*^3^ to 1*10^4^ PFU/ml on day 3 post-infection, which was 1 log lower than in mock vaccinated ferrets (Figure 6). By day 5 post-infection, there was no detectable virus in any COBRA-WIV vaccinated group of ferrets, as well as COBRA-SIV plus R-DOTAP vaccinated ferrets (Figure 6). Low to undetectable viral titers were observed in ferrets vaccinated with COBRA-SIV only or COBRA-SIV plus AddaVax.

**Figure 5.**
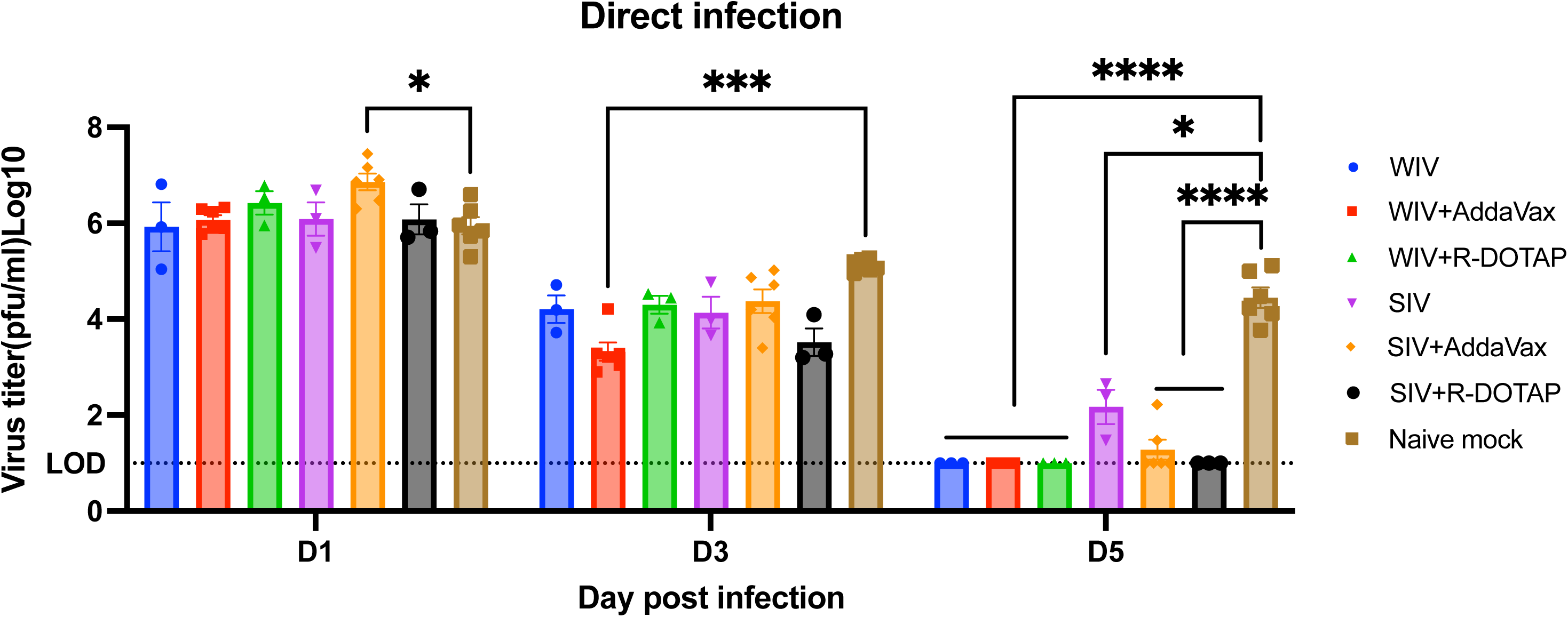
Viral shedding titer in nasal washes collected post-direct infection. The X-axis represents the different time points for collecting the nasal washes. The Y-axis represents the viral titer PFU/ml nasal washes in the Log10 scale with absolute mean values ± SEM. The legend shows the different vaccine groups. A P value of less than 0.05 was defined as statistically significant (*, P < 0.05; **, P < 0.01; ***, P < 0.001; ****, P < 0.0001).

**Figure 6.**
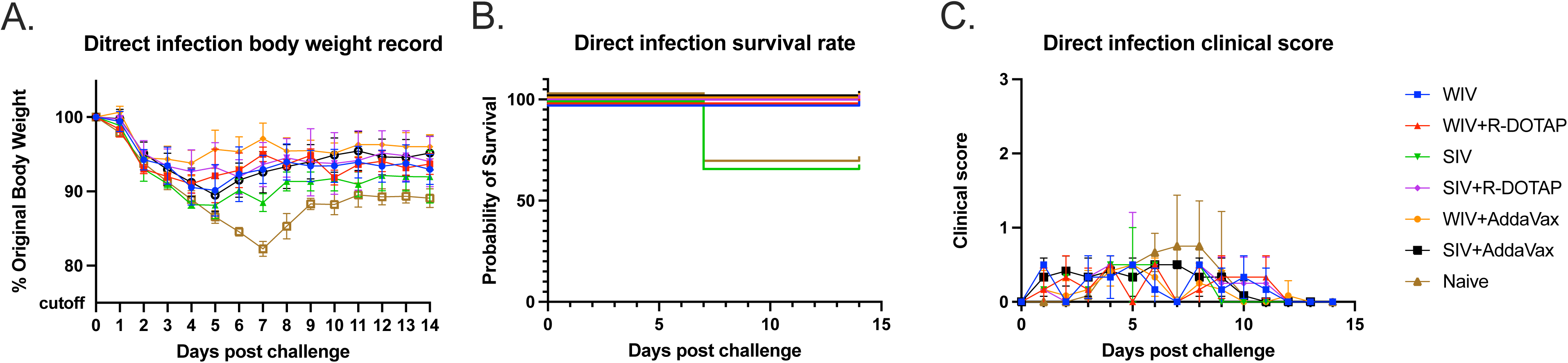
Development of disease post-direct infection. A: body weight loss curve. B: survival rate. C. clinical sores. The legend shows the different vaccine groups with the naïve mock-vaccinated control.

To better assess the relationship between the antibody responses and protection, antibody titer, body weight loss, clinical score, and viral shedding titers were recorded for each individual ferret (Table 2). The results showed that ferrets with higher CA/09-specific antibody titers (HAI>1:40) tended to lose less body weight (<10%) and show fewer clinical symptoms. The viral shedding amount is similar among all ferrets on 1DPI and 3DPI followed by direct infection, but on 5DPI, ferrets with unprotective antibody levels shed more viruses than ferrets with protective antibody levels.

**Table 2.** Individual animal antibody levels, body weight loss, clinical scores, survival days, and viral titers in nasal washes post-infection.^1^.

| Ferret ID | Vaccine | 14 wks post-vax CA/09-specific |  | Infection type | Lowest body weight | Max clinical score | Survival days | Viral shredding (PFU/ml) Log10 |  |  |  |  |
| --- | --- | --- | --- | --- | --- | --- | --- | --- | --- | --- | --- | --- |
|  |  | HAI titer (Log2) | MNA titer |  |  |  |  | 1DPI | 2DPI | 3DPI | 4DPI | 5DPI |
| T1995 | WIV | 5.32 |  | Direct infection | 94.48% | 0.5 | >14DPI | 5.04 |  | 3.72 |  | Undetectable |
| T1996 | WIV | 6.32 |  | Direct infection | 83.37% | 0.5 | >14DPI | 6.82 |  | 4.72 |  | Undetectable |
| T1980 | WIV | 6.32 |  | Direct infection | 92.33% | 0.5 | >14DPI | 5.92 |  | 4.20 |  | Undetectable |
| T1978 | WIV+AddaVax | 2.32 |  | Direct infection | 91.71% | 0.5 | >14DPI | 6.54 |  | 4.45 |  | Undetectable |
| T1979 | WIV+AddaVax | 5.32 |  | Direct infection | 85.41% | 0.5 | >14DPI | 6.78 |  | 4.53 |  | Undetectable |
| T1284 | WIV+AddaVax | 6.32 |  | Direct infection | 97.73% | 0.5 | >14DPI | 5.96 |  | 3.93 |  | Undetectable |
| T372 | WIV+AddaVax | 6.32 |  | Direct infection | 91.71% | 0.5 | >14DPI | 5.89 |  | 4.22 |  | Undetectable |
| T1973 | WIV+AddaVax | 7.32 |  | Direct infection | 95.85% | 0.5 | >14DPI | 6.33 |  | 3.04 |  | Undetectable |
| T1975 | WIV+AddaVax | 5.32 |  | Direct infection | 93.09% | 0.5 | >14DPI | 5.78 |  | 2.90 |  | Undetectable |
| T1281 | WIV+R-DOTAP | 4.32 |  | Direct infection | 89.90% | 0.5 | >14DPI | 6.30 |  | 3.22 |  | Undetectable |
| T127/ T370 | WIV+R-DOTAP | 4.32 |  | Direct infection | 87.19% | 0.5 | >14DPI | 5.90 |  | 3.29 |  | Undetectable |
| T376 | WIV+R-DOTAP | 5.32 |  | Direct infection | 92.86% | 0.5 | >14DPI | 6.24 |  | 3.28 |  | Undetectable |
| T1989 | SIV | 2.32 |  | Direct infection | 84.79% | 1 | >14DPI | 6.69 |  | 4.77 |  | 2.65 |
| T1990 | SIV | 2.32 |  | Direct infection | 88.65% | 0.5 | >14DPI | 6.10 |  | 3.66 |  | 2.40 |
| T1976 | SIV | 4.32 |  | Direct infection | 86.34% | 0.5 | 7DPI | 5.49 |  | 3.98 |  | 1.48 |
| T1984/403 | SIV+AddaVax | 2.32 |  | Direct infection | 92.14% | 0.5 | >14DPI | 7.45 |  | 5.03 |  | 2.22 |
| T1985 | SIV+AddaVax | 4.32 |  | Direct infection | 89.73% | 0.5 | >14DPI | 6.30 |  | 3.41 |  | Undetectable |
| T373 | SIV+AddaVax | 3.32 |  | Direct infection | 85.20% | 0.5 | >14DPI | 7.16 |  | 4.87 |  | Undetectable |
| T395 | SIV+AddaVax | 2.32 |  | Direct infection | 81.36% | 0.5 | >14DPI | 6.48 |  | 4.72 |  | 1.48 |
| T1974 | SIV+AddaVax | 6.32 |  | Direct infection | 85.05% | 0.5 | >14DPI | 6.91 |  | 4.20 |  | Undetectable |
| T1988 | SIV+AddaVax | 3.32 |  | Direct infection | 94.89% | 0.5 | >14DPI | 6.87 |  | 4.04 |  | Undetectable |
| T377 | SIV+R-DOTAP | 7.32 |  | Direct infection | 98.12% | 0.5 | >14DPI | 5.84 |  | 3.27 |  | Undetectable |
| T1983/402 | SIV+R-DOTAP | 5.32 |  | Direct infection | 94.26% | 0.5 | >14DPI | 5.71 |  | 3.20 |  | Undetectable |
| T1997 | SIV+R-DOTAP | 4.32 |  | Direct infection | 86.84% | 1 | >14DPI | 6.71 |  | 4.10 |  | Undetectable |
| T1982 | WIV | 4.32 |  | Contact infection | 93.01% | 0.5 | >14DPI |  | 3.58 |  | 5.63 |  |
| T1986 | WIV | 2.32 |  | Contact infection | 95.24% | 0.5 | >14DPI |  | 2.85 |  | 5.07 |  |
| T1987 | WIV | 4.32 |  | Contact infection | 97.07% | 0.5 | >14DPI |  | Undetectable |  | 4.98 |  |
| T1981 | WIV+R-DOTAP | 6.32 |  | Contact infection | 96.80% | 0.5 | >14DPI |  | 4.82 |  | 4.43 |  |
| T1993 | WIV+R-DOTAP | 5.32 |  | Contact infection | 91.97% | 0.5 | >14DPI |  | 4.73 |  | 4.95 |  |
| T1994 | WIV+R-DOTAP | 2.32 |  | Contact infection | 93.84% | 0.5 | >14DPI |  | 2.70 |  | 5.41 |  |
| T1977 | SIV | 3.32 |  | Contact infection | 91.58% | 0.5 | >14DPI |  | 3.04 |  | 6.20 |  |
| T1991 | SIV | 2.32 |  | Contact infection | 90.48% | 0.5 | >14DPI |  | 2.40 |  | 5.19 |  |
| T1992 | SIV | 2.32 |  | Contact infection | 89.52% | 0.5 | >14DPI |  | 5.21 |  | 5.57 |  |
| T863 | SIV+R-DOTAP | 7.32 |  | Contact infection | 93.82% | 0.5 | >14DPI |  | 2.97 |  | 5.08 |  |
| T1269 | SIV+R-DOTAP | 4.32 |  | Contact infection | 82.12% | 0.5 | >14DPI |  | 5.29 |  | 4.06 |  |
| T1280 | SIV+R-DOTAP | 4.32 |  | Contact infection | 93.22% | 0.5 | >14DPI |  | 5.89 |  | 5.26 |  |
| T1469 | N/A | 2.32 |  | Direct infection | 83.33% | 0.5 | >14DPI | 5.96 |  | 5.20 |  | 4.12 |
| T300 | N/A | 2.32 |  | Direct infection | 83.73% | 0.5 | >14DPI | 5.30 |  | 5.28 |  | 3.77 |
| T1533 | N/A | 2.32 |  | Direct infection | 79.22% | 2 | 7DPI | 5.73 |  | 4.95 |  | 5.11 |
| T3958 | N/A | 2.32 |  | Direct infection | 82.42% | 0.5 | >14DPI | 6.60 |  | 5.06 |  | 4.23 |
| T3966 | N/A | 2.32 |  | Direct infection | 79.48% | 2 | 7DPI | 5.85 |  | 5.03 |  | 4.48 |
| T3967 | N/A | 2.32 |  | Direct infection | 83.59% | 0.5 | >14DPI | 6.26 |  | 5.24 |  | 5.01 |
| T1468 | N/A | 2.32 |  | Contact infection | 85.41% | 1 | >14DPI |  | 6.26 |  | 5.88 |  |
| T1470 | N/A | 2.32 |  | Contact infection | 79.70% | 2 | >14DPI |  | 5.48 |  | 5.01 |  |
| T1542 | N/A | 2.32 |  | Contact infection | 81.31% | 1 | >14DPI |  | 6.10 |  | 5.48 |  |
| T3961 | N/A | 2.32 |  | Contact infection | 87.17% | 0.5 | >14DPI |  | 4.04 |  | 5.83 |  |
| T3971 | N/A | 2.32 |  | Contact infection | 91.23% | 0.5 | >14DPI |  | Undetectable |  | 6.82 |  |
| T3969 | N/A | 2.32 |  | Contact infection | 84.26% | 1 | >14DPI |  | 5.98 |  | 5.88 |  |

<sup>1</sup> HAI, hemagglutination inhibition; MN, microneutralization assay.

To determine if vaccination with COBRA-IIV vaccines could reduce or block respiratory viral transmission, vaccinated ferrets (transmitters) were challenged and then housed with a new, unvaccinated naïve ferret (receiver) for 14 days (Figure 7). Receiver ferrets, housed with vaccinated transmitter ferrets, had nasal wash viral titers that ranged on average between 1*10^5^ to 1*10^6^ PFU/ml regardless which vaccinated ferret was co-housed with the naïve receiver ferret (Figure 7B). The naïve receiver ferrets when housed with vaccinated transmitter ferrets lost significant body weight, with ferrets on average losing between 12-20% body weight by day 8-9 post-infection before their weights plateaued or slightly rose by day 14 post-infection (Figure 8A). Two of the 3 receiver ferrets housed with COBRA-WIV only vaccinated transmitter ferrets and 1 of the 3 receiver ferrets housed with COBRA-WIV plus R-DOTAP vaccinated transmitter ferrets succumbed to infection, while all other receiver ferrets survived (Figure 8B). These two groups’ clinical scores peaked from 7-10 days post-infection with severe lethargy and sharp body weight loss (>20%), which was even higher than those housed with mock-vaccinated transmitter ferrets (Figure 8C).

**Figure 7.**
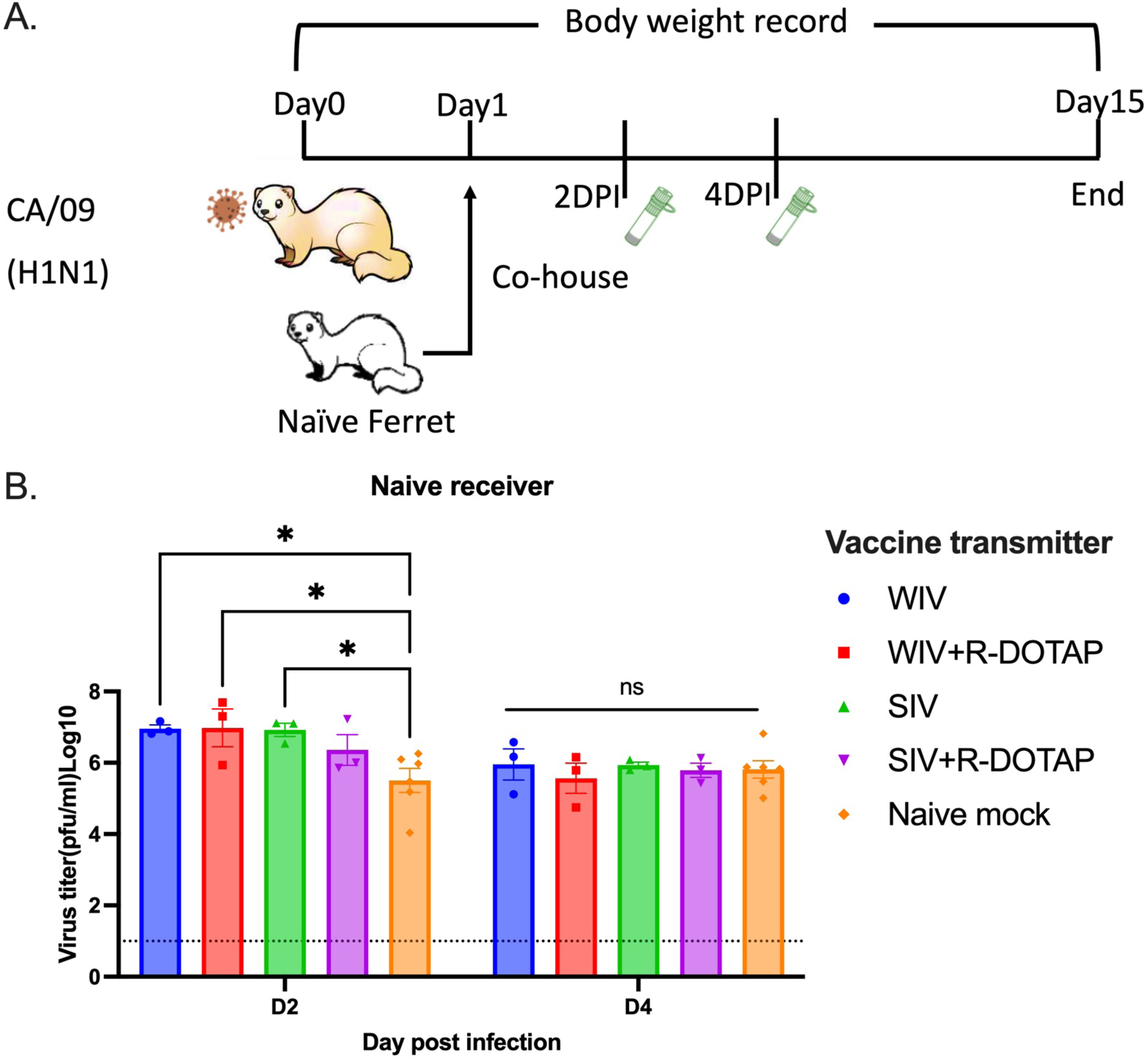
Transmission study from vaccinated transmitters to naive receivers. A. experimental design. Vaccinated transmitters were intranasally infected by CA/09 (1*10^6^PFU/ferret) on day 0. One day later, the naïve receivers were paired with the vaccinated transmitters and they were co-housed for 14 days. The transmission from naïve transmitters to naïve receivers functioned as the mock control in this study. The nasal washes from the receivers were collected on day 3 and 5, which were considered 2DPI and 4DPI for receivers. Body weight loss, clinical scores, and survival rate were closely monitored for 14 days since co-housing. B. Viral titers in nasal washes collected from receivers at 2DPI and 4DPI. The X-axis represents the different time points for collecting the nasal washes. The Y-axis represents the viral titer PFU/ml nasal washes in the Log10 scale with absolute mean values ± SEM. The legend shows the vaccine that the transmitters got. A P value of less than 0.05 was defined as statistically significant (*, P < 0.05; **, P < 0.01; ***, P < 0.001; ****, P < 0.0001).

**Figure 8.**
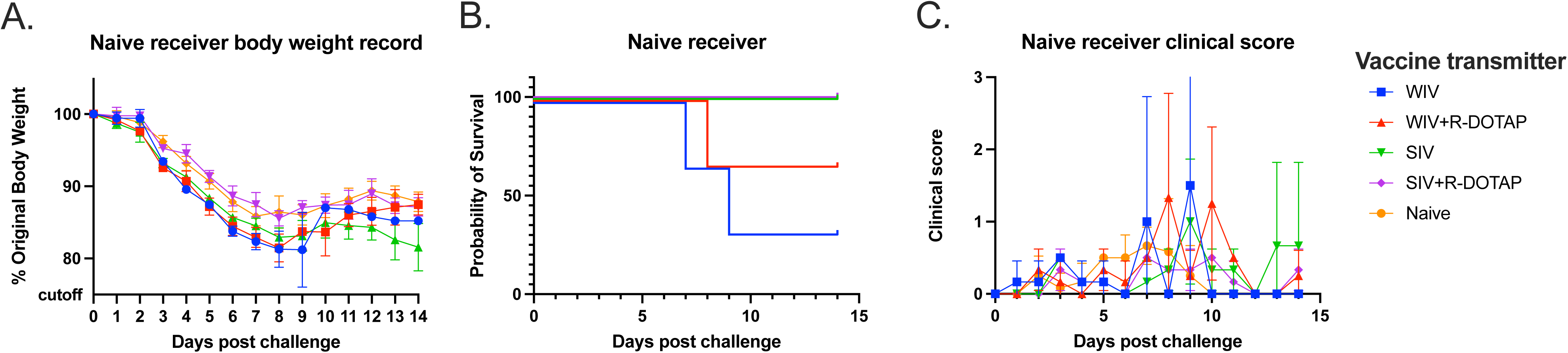
Development of disease post-contact infection in naïve receivers. A: body weight loss curve. B: survival rate. C. clinical sores. The legend shows the different vaccine groups with the naïve mock-vaccinated control.

### Contact infection induced less body weight loss and delayed viral shedding than direct infection in COBRA-IIV vaccinated ferrets

To determine if ferret-to-ferret transmission between animals can be reduced or blocked if the receiver ferret was previously vaccinated, new naïve ferrets were infected with CA/09 and then co-housed with a vaccinated ferret (Figure 9). Mock vaccinated receiver ferrets had ∼10^4.5^ PFU/ml of virus in their nasal washes following 2 days of co-housing with a naïve CA/09 infected transmitter ferret (Figure 9A). These titers rose to ∼10^6^ PFU/ml at day 4 post-infection. Ferrets vaccinated with the COBRA-WIV vaccine only had the lowest viral nasal wash titers (∼10^2.2^) at day 2 post-infection vaccination (Figure 9B). All other vaccinated ferrets had slightly higher, but not statistically different, nasal wash viral titers than COBRA-WIV vaccine only vaccinated ferrets. By day 4 post-infection, all ferrets had similar high nasal wash viral titers as unvaccinated ferrets (between 10^5^ and 10^6^ PFU/ml), regardless of the vaccine administered to the receiver ferrets. Ferrets vaccinated with the COBRA-WIV vaccine with or without R-DOTAP lost between 2-5% of original body weight by day 6 post-infection (Figure 9C). In contrast, ferrets vaccinated with COBRA-SIV vaccines had similar weight loss as unvaccinated ferrets losing ∼10% of their original body weight by day 6 post-infection. The COBRA-SIV vaccinated ferrets began to recover weight over the next 7 days, whereas the unvaccinated ferrets lost between 13-15% of original body weight and plateaued without recovering body weight over the next 7 days of observation (Figure 9C). The individual data shows that even with no detectable CA/09-specific antibodies, some vaccinated ferrets were still protected from the lethal CA/09 challenge and lost <10% of their original body weight (Table 2), which underscores the activation of cellular immunity in protection.

**Figure 9.**
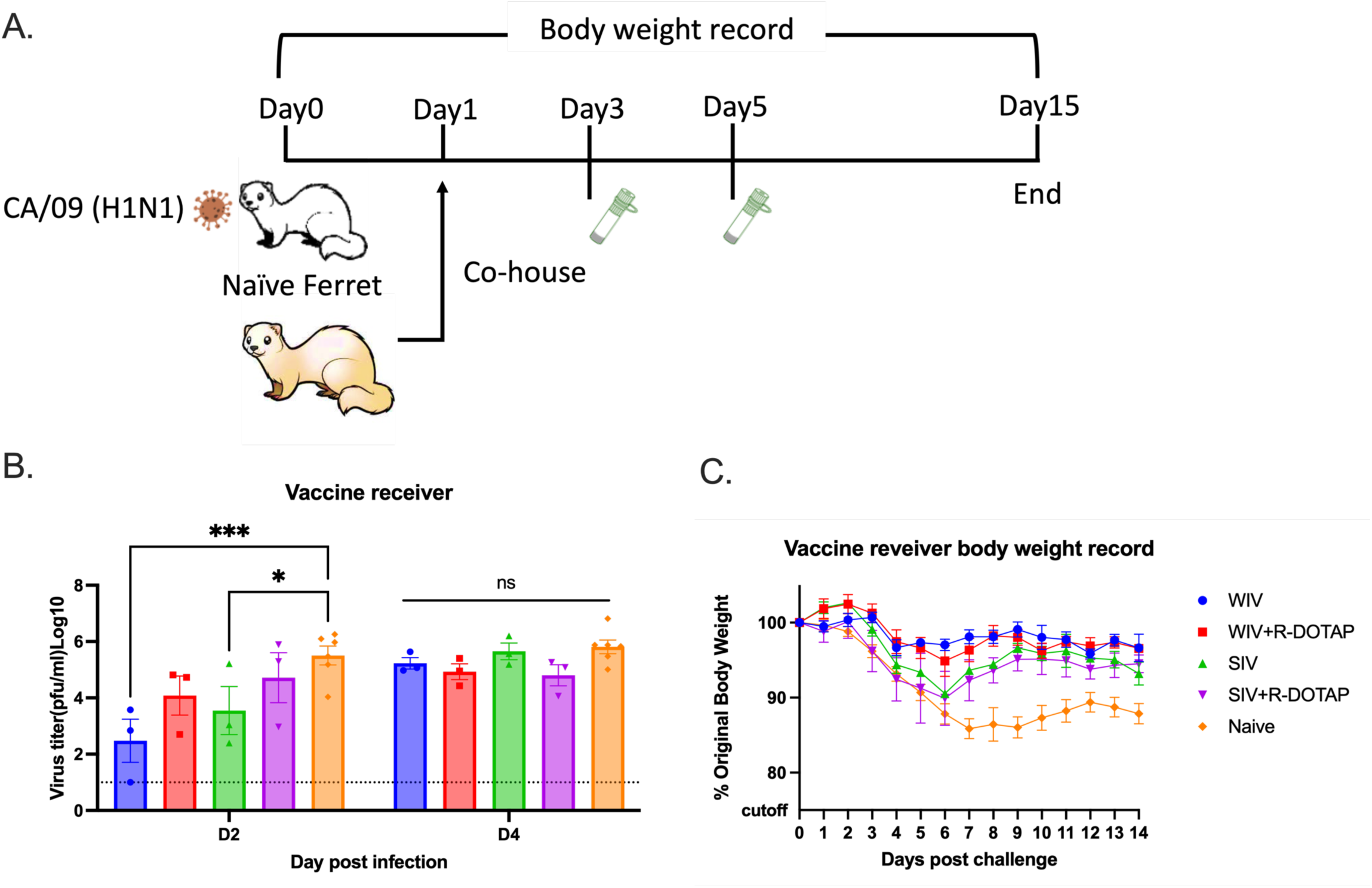
Transmission study from naïve transmitters to vaccinated receivers. A. experimental design. Naïve ferrets were intranasally infected by CA/09 (1*10^6^PFU/ferret) on day 0. One day later, the naïve transmitters were paired with the vaccinated receivers and they were co-housed for 14 days. The transmission from naïve transmitters to naïve receivers functioned as the mock control in this study. The nasal washes from the receivers were collected on day 3 and 5, which were considered 2DPI and 4DPI for receivers. Body weight loss, clinical scores, and survival rate were closely monitored for 14 days since co-housing. B. Viral titers in nasal washes collected from receivers at 2DPI and 4DPI. The X-axis represents the different time points for collecting the nasal washes. The Y-axis represents the viral titer PFU/ml nasal washes in the Log10 scale with absolute mean values ± SEM. C. body weight loss for receivers post-contact infection. The legend shows the different vaccine groups. A P value of less than 0.05 was defined as statistically significant (*, P < 0.05; **, P < 0.01; ***, P < 0.001; ****, P < 0.0001).

Compared to directly infected ferrets, the average body weight loss was less in contact-infected naïve ferrets or ferrets vaccinated with WIV, WIV plus R-DOTAP, and SIV, but similar for ferrets vaccinated with SIV plus R-DOTAP (Figure S3). There was a gap in body weight loss clearly shown between direct infection and contact infection for WIV plus R-DOTAP and WIV vaccinated ferrets, even though no statistical difference was shown (Figure S3). Besides, the naïve ferrets only showed mild clinical signs, such as sneezing and nasal discharge, after the contact infection, but showed some lethargy after the direct infection.

## Discussion

In 2018, the U.S. National Institute of Allergy and Infectious Disease (NIAID) launched a strategy to develop a universal influenza vaccine to ameliorate the global influenza burden (25). To meet this urgent goal, the COBRA methodology used to generate HA and NA and was applied to various vaccine formats for preclinical studies to test their effectiveness (15, 26, 27). In this study, pre-immune ferrets and transmission models were used to determine the vaccine effectiveness of inactivated influenza virus vaccines expressing COBRA HA proteins. The COBRA-IIV vaccines elicited broadly protective neutralizing antibodies against multiple H1N1 and H3N2 influenza virus strains following a single vaccination in pre-immune ferrets.

COBRA-IIV vaccines were effective in both immunologically naïve and pre-immune mice (H. Shi, X. Zhang, P. Ge, V. Meliopoulos, P. Freiden, B. Livingston, S. Schultz-Cherry, T.M. Ross, submitted for publication). In this study, ferrets were used to further assess these vaccine candidates because the mouse-influenza model has some limitations. Typically, mice are resistant to infection by most human influenza strains (28), therefore, mouse-adapted strains are often required for viral challenge (29). Moreover, mice do not show similar clinical symptoms as humans and they poorly transmit viruses between mice (30). Ferrets, another primary animal model for influenza vaccine research, are naturally susceptible to influenza viruses and develop similar clinical signs as humans, such as sneezing, nasal discharge, fever, and lethargy (31). More importantly, the human influenza strains transmit efficiently among ferrets, which makes ferrets ideal for studying viral transmission (32–34).

The WIV vaccine was first used as the influenza vaccine in the 1940s and showed effective protection against influenza virus infection in humans (17). The WIV vaccine maintains conformational structures of viral surface glycoproteins and internal gene products (35). However, the strong immunogenicity was associated with inflammation at the site of administration resulted in limited human use in the U.S. (36). The SIV vaccines with improved safety has been used as a commercial seasonal influenza virus vaccine for decades and is safe and effective in most populations (18). However, the SIV vaccine is not as effective in children and the elderly (21, 37, 38). The reduced vaccine effectiveness elicited by SIV vaccines compared to WIV vaccines was also observed in this study. The COBRA-WIV elicited antibodies with broad HAI activity (9 out of 11 tested strains), whereas the COBRA-SIV elicited HAI activity at >1:40 against 3 out of 11 strains in the HAI panel (Figure 2). In addition, the COBRA-WIV stimulated significantly more HA stem-binding antibodies than the COBRA-SIV (Figure 4). With the optimized inactivation bioprocess and purification methods, the adverse effects of WIV vaccines can be diminished (19). In this study, no vaccination-associated reactions were observed after administrating the COBRA-WIV vaccines to ferrets. To enhance the vaccine effectiveness of SIV vaccines, a higher dose of vaccine or addition of an adjuvant has been used in commercial split-inactivated influenza vaccines for the elderly (39). Similarly, when the COBRA-SIV vaccines were mixed with R-DOTAP adjuvant, the antibody breadth and protection against viral infection were improved. Meanwhile, the AddaVax did not significantly enhance the vaccine effectiveness compared to unadjuvanted vaccine. Therefore, the utilization of COBRA-SIV may require an adjuvant if tested in the clinic.

R-DOTAP is designed to attach to the negatively charged cell membrane and deliver the peptide or protein antigens into the host cells, which stimulates cellular immunity and cross-presentation (40). R-DOTAP adjuvanted COBRA rHA vaccines elicit antibodies with wider HAI breadth and stimulated more multifunctional T cells (CD8+ and CD4+ T cells) than unadjuvanted COBRA rHA vaccines (24). The addition of R-DOTAP to COBRA-SIV improved the antibody activity spectrum (Figure 2F) with only one vaccination and induced more stem-binding antibodies than unadjuvanted COBRA-SIV or AddaVax adjuvanted COBRA-SIV (Figure 4B). Compared to AddaVax, R-DOTAP is likely a better adjuvant for COBRA-SIV formulation, since it further improved the breadth HAI activity elicited by COBRA-SIV (Figure 2 E and F), and was more effective at suppressing viral replication following infection in ferrets (Figure 5). Furthermore, the R-DOTAP stimulates more durable, poly-functional CD4+ and CD8+ T cells than with AddaVax and these T cell receptors (TCRs) also recognized by more distinct influenza peptides (24). Future studies will evaluate the cellular immunity elicited by COBRA-IIV vaccines and the map TCR usage in order to optimize the vaccine formulation.

However, the use of either adjuvant had a few positive or even negative effects on the COBRA-WIV in eliciting stem-binding antibodies (Figure 4B), which could result from the self-adjuvant effects of WIV (41, 42). Since the COBRA-WIV stimulated robust immune responses by itself, the addition of an adjuvant did not introduce significant improvement. A similar result was also noticed in the mouse study that the COBRA-WIV and COBRA-WIV plus AddaVax elicited comparable HAI titers and breadth in naïve mice after two vaccinations (H. Shi, X. Zhang, P. Ge, V. Meliopoulos, P. Freiden, B. Livingston, S. Schultz-Cherry, T.M. Ross, submitted for publication). However, the addition of an adjuvant sustained the antibody circulation for both COBRA-WIV and COBRA-SIV (Figure S1). At 14 weeks post-vaccination, the COBRA-WIV plus AddaVax and COBRA-SIV plus R-DOTAP maintained a protective HAI level against CA/09, while other formulations all waned below 1:40 (Figure S1).

while all other formulations, the HAI titer waned ≥1:40 (Figure S1). The addition of an adjuvant provided better protection against the lethal challenge by direct infection (Figure 6) or by contact infection (Figure 9 and Figure S2).

To mimic person-to-person transmission patterns in humans, vaccinated ferrets were co-housing with infected ferrets. Sixty-six percent of immunologically naïve ferrets survived challenge when directly infected with CA/09 (H1N1) (Figure 6B). These ferrets lost more weight post-infection compared to ferrets infected by contact transmission. In addition, 100% of ferret survived infection via contact transmission from another ferret. Interestingly, the different infection methods also led to different outcomes using the same vaccine (Figure S3). The COBRA-SIV vaccinated ferrets had higher fatality and more body weight loss than the COBRA-SIV plus R-DOTAP following direct infection (Figure 6A and B). However, after contact infection, these two vaccines elicited similar protective immunity that resulted in improved survival and less weight loss (Figure 9C). This may be a result that the receiver ferrets were exposed to the gradually increasing doses virus during transmission. In contrast, the directly infected ferrets were inoculated with a lethal dose of the virus, which causes an acute infection (43). The infection methods led to different interpretations for the vaccine effectiveness of COBRA-SIV and COBRA-SIV plus R-DOTAP, which emphasizes the importance of model choice in preclinical vaccine studies.

The goal of vaccination is not only to protect the vaccinated individuals against disease and illness but also to reduce shedding and transmission to non-vaccinated people. This is known as herd immunity. The COBRA-IIV vaccines provided protection for over 14 weeks, however, vaccinated ferrets consistently shed virus that was similar to viral titers observed in mock-vaccinated ferrets at 1-day post-infection (Figure 5). This phenomenon was also observed in the previous ferret and pig studies (33, 44). One possible explanation is that the COBRA-IIV vaccines did not elicit sufficient antibodies to neutralize all viruses. As shown in Figure S1, the CA/09-specific antibodies in most ferrets dropped below the protective level. Therefore, those viruses still infected the vaccinated ferrets and replicated rapidly in their respiratory tract. However, significantly faster elimination of virus was observed in the vaccinated ferrets than in the naïve control, which indicates an overall reduced viral shedding. The CA/09 is a highly transmissible virus via aerosols or droplets containing only a low viral load (45) and the naïve receivers were exposed to the transmitters for 14 days, which led to infection in the naïve receivers (Figure 8B).

Even though COBRA-IIV vaccines did not block transmission in this study, the period of viral shedding in vaccinated ferrets was shorter than in naïve ferrets (Figure 5). All vaccinated ferrets did not shed virus or shed significantly less viruses than the naïve mock control ferrets at 5 days post-infection (Figure 5), even for ferrets that had HAI antibody levels >1:40 prior to challenge (Figure S1). This may indicate that the immune memory was activated and responded rapidly to eliminate virally infected cells within 5 days. Furthermore, vaccinated ferrets infected through contact had delayed shedding patterns and one ferret (T1987) did not shed any virus at 2 days post-infection (Figure 9B). Both the timing of the exposure and the period of exposure are important factors in determining transmission. The co-housing transmission model does not effectively mimic the public transmission pattern of human model with potentially longer exposure times via droplet, aerosol, and direct contact. However, the primary source of influenza virus transmission in the human population is via droplets or aerosol containing infectious particles with a short exposure time (46). With shorter exposure time, the viral shedding in COBRA IIV-vaccinated ferrets may be blocked. For example, if a naïve ferret is exposed to T1987 at 1-2 days post-infection for a short time and then separated, the ability to transmit virus between animals will be low because no virus was detectable for 2 days post-infection (Figure 7B). Moreover, direct contact introduces more infectious particles than droplets or aerosols (47). Therefore, in future studies, a more sophisticated transmission model in which the receivers are exposed to the infected transmitters without direct contact in a short time period should be explored. In this way, the effectiveness of COBRA-IIV vaccines to block viral transmission from vaccinated transmitters to naïve receivers can be further assessed. Additionally, designing different timing(s) of exposure, especially before the clinical signs appear will provide more details about the contagious dynamics after influenza virus infection (https://www.cdc.gov/flu/about/keyfacts.htm).

In humans, HAI titers induced by vaccination that are ≥1:40 are associated with 50% protection against influenza virus infection (48). In this study, ferrets that received COBRA-WIV and COBRA-SIV plus R-DOTAP maintained a protective HAI titer against CA/09 before challenge, but these ferrets were not protected from the CA/09 challenge, which may be associated with the dose of virus dose used in this study. The different exposure doses of influenza virus lead to different immune responses and consequentially effect vaccine effectiveness (49). Influenza virus dose as low as 1.95*10^3^ viral copies can effectively infect people (50), which is 3 logs lower than the dose (10^6^ infectious particles/ferret) used to infect ferrets in this study. This high challenge dose is to ensure the experimental infection and transmission, which may limit the vaccine induced protective immune responses elicited by COBRA-IIV vaccines. Future studies can address various doses of vaccine and adjuvants, as well as various challenge doses of virus.

In summary, COBRA IIV vaccines elicited broadly reactive antibodies and provided long-term protection with only one dose in pre-immune ferrets. Among all tested formulations, the COBRA-WIV showed the most promising effectiveness, as well as its ease of manufacturing (no extra splitting process) and no need for an adjuvant. Additionally, the COBRA-SIV plus R-DOTAP provided comparable protection against the lethal challenge as the COBRA-WIV. Considering that the current influenza virus vaccines are manufactured in the SIV format, the COBRA-SIV plus R-DOTAP is also a promising candidate for humans. Overall, the COBRA-WIV and COBRA-SIV plus R-DOTAP provided broad and durable protection and reduced viral transmission.

### Materials and methods Vaccine preparation

The recombinant viruses expressing COBRA HA Y2 (H1N1) or J4 (H3N2) were rescued by collaborators in St. Jude Children’s Hospital by using the eight-plasmid reverse transgene system and then inactivated and split by us to generate the COBRA-WIV and COBRA-SIV. This method has been described previously. Briefly, the mixture of 293T (6.25× 10^5 cells/well) and Madin-Darby canine kidney (MDCK) cells (3.2510^5 cells/well) was seeded in 6-well cell culture plates until they achieved 80-90% confluency. The cDNA of each segment of A/Puerto Rico/8/1934 (PR8) was inserted into the plasmid vector pHW2000, except for the segment encoding the HA. It was substituted with the gene encoding the COBRA HAs. Plasmids for virus recovery were prepared meticulously, ensuring each attained a concentration as close to 1mg/ml as feasible. 1μg of each plasmid was added to each well (8μg in total) mixed with 500μL OptiMEM (ThermoFisher, 31985-062) and 16μL TransIT-LT (Mirus, MIR2300), followed by overnight incubation at 37°C with 5% CO2. The subsequent day involved replacing the medium with OptiMEM containing TPCK-Trypsin (Worthington, LS003740). As the days progressed, diligent monitoring for cytopathic effects (CPE) and hemagglutination (HA) was necessary. If required, blind passages were conducted, and the supernatant was stored at -80°C for future use.

To amplify the rescued virus, MDCK cells were prepared and when they achieved 80-90% confluency, the cell culture media was discarded and replaced with the viruses (MOI: 0.01) diluted in 1XMEM (Corning). After 48-72 hours of incubation at 37°C with 5% CO2, the cells were monitored for CPE, and when obvious CPE was observed, the cell culture media was collected and centrifuged at 2000xg and 4°C for 10 minutes to separate the supernatant. The supernatant was carefully collected and mixed with a 0.5M disodium phosphate (DSP) solution in a ratio of 1 part DSP solution to 38 parts supernatant. Next, 2% BPL was added dropwise to the DSP-supernatant mixture in a ratio of 1 part BPL to 38 parts DSP-supernatant solution, resulting in a final BPL concentration of 0.05% (v/v). The mixture was then incubated on ice with intermittent shaking for 30 minutes before being transferred to a 37°C water bath for 2 hours with periodic mixing. Following incubation, the pH of the mixture was adjusted to 7.3-7.4 by using a 7% sodium bicarbonate solution, monitored with pH strips. The resulting antigen are the COBRA-WIV. They were quantified by anti-H1 or anti-H3 western blot and stored at -80°C until further use. Additionally, an inoculation test was carried out to detect the presence of any live viruses, providing further assurance of the antigen’s safety for use in subsequent applications.

To produce the COBRA-SIV, a 1% (v/v) final concentration of Triton X100 was introduced into the WIV solution, followed by incubation at room temperature for one hour. Subsequently, to ensure safety, 0.1g of BioBeads per mL of the mixture was incorporated to eliminate excess Triton X100. The mixture underwent gentle shaking on a shaker at 4°C overnight, after which the liquid component was separated and collected. In this way, the COBRA-SIV was generated. A similar western blot was conducted to quantify the HA content and then the COBRA-SIV was labeled and stored at -80°C for further use.

### Animal, pre-infection, and vaccination

Female Fitch ferrets (Mustela putorius furo) aged 6 to 15 months were obtained from Triple F Farms (Gillett, PA, USA) after de-scenting and spaying procedures. They were resting in the animal facility for one week before any procedures. Blood samples were collected as the baseline for each ferret’s serological status regarding the A/California/07/2009 H1N1 influenza virus. Ferrets were housed in pairs with access to food, water, and enrichment when not undergoing procedures. Anesthesia using vaporized isoflurane was administered before any interventions such as bleeds, vaccination, infection, nasal washes, or euthanasia. Ethical standards, as outlined in the Guide for the Care and Use of Laboratory Animals, Animal Welfare Act, and Biosafety in Microbiological and Biomedical Laboratories (AUP: A2020 11-016-Y1-A6), were strictly followed during all animal procedures. To establish pre-immunity, naïve ferrets were intranasally infected with the Sing/86 and Pan/99 influenza virus (5*10^5^PFU/virus) eight weeks prior to the vaccination. Following this, ferrets (pre-immune or naïve) were vaccinated intramuscularly in the thigh muscle with vaccines (7.5μg/antigen) formulated as described in Table 1. The Addavax adjuvant (InvivoGen, San Diego, CA, USA) or R-DOTAP (PDS, Florham Park, NJ, USA) was mixed in a 1:1 ratio with sterile PBS and antigens. Naive mock-vaccinated groups received 500μL of sterile PBS. Blood samples were collected in BD Vacutainer SST tubes 2-week post-preinfection to confirm the pre-existing antibodies specific to Sing/86 and Pan/99; and 2-, 10-, and 14-week post-vaccination. Serum separation was achieved by centrifugation at 2500 rpm for 10 minutes following 30 minutes of incubation at room temperature. The purified serum was then stored at −20 °C until further analysis. The experimental timeline is shown in Figure 1.

### Viruses

All viral strains employed in this study were sourced from Influenza Reagents Resource (IRR), BEI Resources, the Centers for Disease Control (CDC), or graciously provided by Virapur (San Diego, CA). Those viruses were amplified strictly following the environment of their original stocks. The H1N1 influenza viruses included A/Singapore/6/1986 (Sing/86), A/California/07/2009 (CA/09), A/Brisbane/02/2018 (BS/18), A/Guangdong-Maonan/SWL1536/2019 (GD/19), and A/Victoria/2570/2019 (Vic/19). The H3N2 influenza virus strains included A/Panama/2007/1999 (Pan/99), A/Switzerland/9715293/2013-mouse adapted (SW/13), A/Hong Kong/4801/2014 (HK/14), A/Singapore/IFNIMH-16-0019/2016 (Sing/16), A/Kansas/14/2017 (Kan/17), A/South Australia/34/2019 (SA/19), A/Hong Kong/2671/2019 (HK/19), and A/Tasmania/503/2020 (TAS/20).

### Direct infection

Ferrets were challenged by CA/09 (1*10^6^PFU/ferret/1mL) via intranasal inoculation at week 14. Following infection, animals were closely monitored, with observations conducted twice daily to detect any potential clinical signs. Additionally, their weight was measured once daily until 14 days post-infection (DPI). Nasal washes were conducted on days 1, 3, and 5 DPI utilizing 3 mL of sterile PBS each time.

### Contact infection

The contact infection was designed in a two-way transmission: from the naïve transmitter (NT) to the vaccinated receiver (VR) and from the vaccinated transmitter (VT) to the naïve receiver (NR). NT ferrets were infected by CA/09 (1*10^6^PFU/ferret/1mL) via intranasal inoculation. One day after the infection, the VR ferrets were paired with the NT ferrets. For the other direction of transmission, VT ferrets were infected by CA/09 (1*10^6^PFU/ferret/1mL) via intranasal inoculation and on 1 DPI, NR ferrets were co-housed with the VT ferrets. The transmission from the NT to the NR serves as the mock control. The NT ferrets were infected by CA/09 (1*10^6^PFU/ferret/1mL) via intranasal inoculation. One day after the infection, the NR ferrets were paired with the NT ferrets.

All co-housed ferrets stayed together all the time for the rest of the experiment. Following infection, animals were closely monitored, with observations conducted twice daily to detect any potential clinical signs. Additionally, their weight was measured once daily until 14 days post-infection (DPI). Transmitters’ nasal washes were conducted on 1, 3, and 5 DPI, and the receivers’ nasal washes were collected on 2 and 4 DPI, utilizing 3 mL of sterile PBS each time.

### Hemagglutination inhibition (HAI) assay

The hemagglutination inhibition (HAI) assay is capable of identifying antibodies that specifically bind to the hemagglutinin (HA) head, thereby preventing the attachment of viral HA to sialic acid receptors on red blood cells. This inhibition results in the prevention of hemagglutination, which can be readily observed. To ensure the assay’s specificity, all serum samples underwent treatment with a receptor-destroying enzyme (RDE) (Denka Seiken, Co., Japan). Serum samples were mixed with RDE at a 1:3 ratio and incubated at 37°C overnight. Subsequently, the mixture was subjected to heat inactivation at 56°C for 45 minutes, followed by the addition of phosphate-buffered saline (PBS) to achieve a final ratio of serum: RDE: PBS of 1:3:6. The RDE-treated serum samples were then subjected to serial dilution in PBS within a 96-well v-bottom plate. For H1N1 viruses, turkey red blood cells (TRBC) sourced from Lampire Biologicals, Pipersville, PA, USA, were diluted in PBS to a final concentration of 0.8%. Each virus was adjusted to a concentration of 8 hemagglutination units (HAU)/50μL, and an equal volume of virus was added to the serum plate. Following a 20-minute incubation period, 0.8% TRBC was added to each well, and the plates were gently tapped to ensure thorough mixing before being incubated at room temperature (RT) for 30 minutes. Conversely, for H3N2 viruses, guinea pig red blood cells (GPRBC) from Lampire Biologicals, Pipersville, PA, were diluted in PBS to a concentration of 0.75%. Similarly, each virus was adjusted to a concentration of 8HAU/50μL in the presence of 20nM Oseltamivir, and an equal volume of the virus was added to the serum plate. After a 30-minute incubation period, 0.75% GPRBC was added to each well, followed by gentle mixing and incubation at RT for 1 hour. HAI titers were determined based on the reciprocal dilution of the last well where no hemagglutination was observed. According to guidelines provided by the World Health Organization (WHO), HAI titers exceeding 1:40 are considered indicative of a protective level (Use, 2016).

### Microneutralization assay

The H1N1virus strains CA/09 and Vic/19 and H3N2 strains Sing/16. KS/17, HK/19, SA/19, and TAS/20, with a final concentration of a 100X tissue infectious dosage 50 (TCID50) per 50μL, underwent incubation with diluted mice serum samples in two-fold serial dilution, ranging from 1:10 to 1:1280, within the virus diluent. One hour of incubation at 37°C, 5% CO2 was conducted to allow the neutralization between the antibodies and viruses. Subsequently, the MDCK cells with an adjusted concentration of about 3*10^5^ cells/mL were added to the mixtures of the virus and serum and allowed to an overnight incubation at 37°C, 5% CO2 for viral growth. Following this incubation period, the cell medium was discarded and fixed with 80% Acetone, followed by 3X wash with wash buffer. 100 μl of the primary antibody Rabbit-anti-influenza-A-NP Polyclonal Antibody (Thermo Fisher, PA5-81661) was diluted 1:2000 and added to the fixed cells followed by a one-hour incubation at room temperature. After 3X wash with wash butter, 100 μl of the secondary antibody Goat-anti-rabbit IgG (H+L) HRP (Thermo Fisher, 31460) 1:2000 diluted was added and subjected to a one-hour incubation at room temperature. Upon completion of the incubation period, the plates were washed 5 times, and 100 μl of o-phenylenediamine dihydrochloride (OPD) (Sigma, cat. no P8287) was added, initiating an enzyme reaction. This reaction was stopped after 3 minutes with the addition of 100 μl of 2N sulfuric acid. Quantification of the reaction was performed by measuring the optical density (OD) at a wavelength of 492 nm. The neutralization (NT) titers were determined as the reciprocal of the serum dilution yielding a reduction in infected cells of ≥ 50% compared to the virus control.

### Enzyme-linked immunosorbent assay (ELISA)

ELISA assays were performed to detect the level of IgG against various antigens, following established protocols (Huang et al., 2021). Immulon 4HBX plates (Thermo Fisher Scientific, Waltham, MA, USA) were coated with a solution containing Y2 rHA, J4 rHA, cH6/1, or cH7/3 at a concentration of 1μg/mL. After overnight incubation in a humidified chamber at 4°C, plates were blocked with 200μL of blocking buffer per well, and incubated at 37°C for 90 minutes. Concurrently, serum samples were diluted in blocking buffer using a 3-fold serial dilution starting from 1:500. Following blocking buffer removal, 100μL of diluted serum was added to each well. After a 1-hour incubation at 37°C, a secondary antibody, biotinylated goat anti-ferret IgG (Sigma-Aldrich, St. Louis, MO), diluted in 1:4000, was added to each well (100μL/well) and incubated at 37°C for 1 hour. Subsequently, 50 μL of ABTS substrate (VWR Corporation) was added to each well and incubated at 37°C for 15 minutes. The reaction was stopped by adding 50 μL of 1% SDS to each well. Optical density (OD 414) values were measured using a spectrophotometer (PowerWave XS, BioTek) at a wavelength of 414 nm.

### Plaque assay

A plaque assay was conducted according to previously established protocols (Allen et al., 2018; Allen & Ross, 2018; Allen & Ross, 2022; Carter et al., 2017). Briefly, MDCK cells were seeded at a density of 5 × 10^5 cells/well in 6-well plates in cell growth media two days prior to the assay initiation. The assay commenced when cell confluence exceeded 90%. Nash washes were subjected to 10-fold serial dilution ranging from 10^0 to 10^-5 in Dulbecco’s modified Eagle medium (DMEM). These diluted samples were sequentially added to individual wells of 6-well plates and incubated at 37°C for 1 hour with gentle rocking every 15 minutes. Following virus attachment, the media were aspirated, and the plates were washed twice with DMEM to remove unbound viruses. Subsequently, 2 mL of 0.8% agarose (Cambrex, East Rutherford, NJ, USA) was added to each well, and the plates were further incubated at 37°C with 5% CO2 for 72 hours. After incubation, the agarose overlay was carefully removed, and the cell monolayers were fixed using a 10% formalin solution, followed by staining with 1% crystal violet (Thermo Fisher). The plates were then rinsed with tap water and allowed to air dry. The number of plaques on each plate was counted, and the plaque-forming units (PFU) per milliliter for each virus were calculated.

### Statistical analysis

Differences in weight loss, dynamic antibody titers against challenge strains, and lung viral titers were analyzed by two-way ANOVA. The differences in HAI titer and IgG titers were analyzed by one-way ANOVA. Statistical significance was defined as a p-value of 0.05. Graphs and statistical analyses were done using GraphPad Prism software.

## Acknowledgment

We would like to thank James Allen, Ivette Nuñez, and Ying Huang for designing COBRA HA vaccines and Spencer Pierce for purifying the HA antigens. We thank Naoko Uno technical assistance. We also thank the University of Georgia Animal Resource staff, technicians, and veterinarians for their excellent animal care. We also appreciate Benjamin Chadwick for proofreading and language editing for this paper. We acknowledge BEI Resources and the International Reagent Resource for providing influenza viruses used in this study. This project has been funded as part of the Collaborative Influenza Vaccine Innovations Centers (CIVICs) by the National Institute of Allergy and Infectious Diseases, a component of the NIH, Department of Health and Human Services, under contract 75N93019C00052. TMR is also supported in part as a Georgia Eminent Scholar by the Georgia Research Alliance, GRA-001.

